# Deletion of two genes improves heterologous secretion of fungal lignin-degrading heme peroxidases in *S. cerevisiae*

**DOI:** 10.1101/2021.11.27.470220

**Authors:** Nikita A. Khlystov

## Abstract

Efficient, large-scale heterologous production of enzymes is a crucial component of the biomass valorization industry. Whereas cellulose utilization has been successful in applications such as bioethanol, its counterpart lignin remains significantly underutilized despite being an abundant potential source of aromatic commodity chemicals. Fungal lignin-degrading heme peroxidases are thought to be the major agents responsible for lignin depolymerization in nature, but their large-scale production remains inaccessible due to the genetic intractability of basidiomycete fungi and the challenges in the heterologous production of these enzymes. In this study, we employ a strain engineering approach based on functional genomics to identify mutants of the model yeast *Saccharomyces cerevisiae* with enhanced heterologous production of lignin-degrading heme peroxidases. We show that our screening method coupling an activity-based readout with fluorescence-assisted cell sorting enables identification of two single null mutants of *S. cerevisiae, pmt2* and *cyt2*, with up to 11-fold improved secretion of a versatile peroxidase from the lignin-degrading fungus *Pleurotus eryngii*. We demonstrate that the double deletion strain *pmt2cyt2* displays positive epistasis, improving and even enabling production of members from all three classes of lignin-degrading fungal peroxidases. We anticipate that these mutant strains will be broadly applicable for improved heterologous production of this biotechnologically important class of enzymes.

## Introduction

Heterologous production of proteins remains a significant bottleneck in biotechnology despite decades of research^1,2^. Heterologous secretion is particularly important for the production of biotechnologically and medically relevant proteins such as enzymes and antibodies, representing multi-billion-dollar industries^3^. Factors governing successful secretion of foreign proteins are however still poorly understood despite existing significant knowledge about the secretory pathway of common eukaryotic heterologous hosts^2,4^. Engineering efficient secretion in these hosts is a long-sought goal with potentially far-reaching implications from pharmaceutical development to biomass valorization^5–7^. A common approach of improving heterologous secretion involves directed evolution campaigns aimed at identifying protein sequence variants that enable greater levels of production^8^. This approach however is marred by a strong dependence on the screening method involved, often resulting in unintentional and sometimes undesirable alteration to molecular properties of the protein target. This holds especially true for enzymes having native substrates that are challenging to employ in high-throughput screening and where small-molecule proxies are instead employed, such as is the case for lignin-degrading enzymes. In these cases, evolution directed at a proxy substrate may not confer enhanced enzyme properties towards the actual substrate of interest. Engineering approaches that maintain native enzyme catalytic properties rely on judicious selection of promoters and expression cassettes^9^ as well as well-tolerated endoplasmic reticulum (ER) signal peptides^10^ to maximize enzyme yield.

Modification of the protein sequence to accomplish efficient secretion requires individual development of variants for each protein of interest, an inherently time-consuming process. Optimization of the genetic over-expression platform is likewise a constrained approach limited by the native capacity of the host’s secretory pathway. Developing a super-secreting heterologous host would circumvent both of these long-standing issues. With the abundance of genetic tools and wealth of knowledge available, *Saccharomyces cerevisiae* presents an attractive candidate for a proof-of-concept super-secreting platform. The secretory pathway of this yeast has been extensively characterized and the secretion of numerous foreign proteins has been studied in this host^11^. Functional genomic screens as well as targeted gene editing approaches have been successfully employed in engineering improved secretion of heterologous proteins. Gene deletion and cDNA overexpression libraries have revealed that modulating expression of native secretory components can result in enhanced secretion^4,12–18^. Reverse metabolic engineering screens have demonstrated that whole-cell random mutagenesis by UV is another powerful approach in developing strain backgrounds with improved secretion^19,20^. Biotechnologically relevant foreign proteins such as antibodies, T-cell receptors, cellulases, and amylases have all been shown to have improved secretion in the context of these engineered strains. However, these strategies suffer from low-throughput screening methods and expensive whole-genome sequencing to identify strain modifications resulting in secretion enhancement.

In this study, we sought to develop a screening strategy that would help overcome the challenges of heterologous production for a class of enzymes important for biomass valorization, fungal lignin-degrading heme peroxidases. Lignin is an abundant yet underutilized renewable biopolymer derived from woody biomass that could serve as a source for commodity chemicals and materials such as vanillin and phenolic resins that are currently unsustainably derived from petroleum^21,22^. Fungal basidiomycete species are well-known voracious degraders of lignin in nature but remain genetically intractable and difficult to cultivate^23,24^. Engineering a synthetic biochemical route to lignin valorization would enable extraction of valuable chemical products but requires heterologous production of lignin-degrading peroxidases, a major bottleneck in the field today^23^. Since lignin is a bulky, insoluble polymer, the use of directed evolution to develop highly secreted enzyme variants requires small molecule substrate proxies^25,26^. Because these substrates only partially approximate lignin’s physiochemical properties, directed evolution of these peroxidases would likely to alter and even reduce their native lignin-degrading properties. Accordingly, we hypothesized that a strain engineering approach would preserve these properties while improving secretion. We describe a high-throughput method for screening peroxidase-producing *S. cerevisiae* strains from a homozygous single null-mutant deletion collection using fluorescence-assisted cell sorting (FACS). We quantitatively assess the impact of each gene deletion, show individual validation of mutant strains, and identify two single null mutants with significantly improved secretion. We show that a double-deletion strain displays further improvement in secretion, indicating positive epistasis for these two genes. Finally, we demonstrate that the engineered strains have increased secretion of members of all three classes of fungal lignin-degrading peroxidases, indicating that the improved strains are broadly applicable for heterologous production of this class of enzymes.

## Results

Since most lignin-degrading peroxidases have limited heterologous secretion in yeast, we focused on horseradish peroxidase (AR-*hrp*) as a model heme peroxidase with structural similarity to lignin-degrading peroxidases^27,28^ and detectable secretion in wildtype strains of *S. cerevisiae*^29^. In order to rapidly screen strains for secretion ability, we chose yeast surface display as a reliable proxy for secretion^30^ to enable pooled library screening by FACS. We adapted previously described labeling methods^31–33^ involving biotinyl tyramide to screen based on both enzyme production as well as activity, ensuring that the enzyme is properly folded and functional in yeast strains having greater levels of protein display.

We operated in the homozygous deletion collection background^34^ containing barcoded knockouts of each non-essential gene in yeast. We cloned a galactose-inducible vector that allowed plasmid-based expression of both the AGA1 scaffold protein and AGA2 fusion protein for surface display of horseradish peroxidase^35,36^. After binning based on the presence or absence of displayed functional enzyme (“productive” versus “non-productive” cells), we employed a barcode counting approach^37^ to quantify the abundance of each deletion strain in the two bins by next-generation sequencing. We determined impact on secretion using the log fold change in abundance between the two bins. 546 strains showed statistically significant improvement in surface display (log fold change > 2, *p*-value < 1e-10), of which 12 were manually selected for subsequent characterization based on ontological relation to the secretory pathway.

We compared production of horseradish peroxidase in these 12 mutant strains to that in a silent deletion strain (Δ*ho*). Even with a growth defect resulting from an inability to utilize nonfermentable carbon sources (**Figure 1**), BY4743Δ*cyt2*, representing deletion of cytochrome c1 heme lyase, showed significant improvement in horseradish peroxidase secretion (3.38-fold greater than wild-type after OD_600_ normalization), especially in the absence of heme supplementation in culture media (9.10-fold) (**Figure 2**). Without heme and calcium media supplementation, BY4743Δ*qdr2* and BY4743Δ*pom152* also showed statistically significant enhancement in secretion (1.75- and 1.31-fold, respectively). We also tested the heterologous production of a versatile peroxidase from the lignin-degrading fungus *Pleurotus eryngii* (PE-*vpl2*) with low levels of secretion in *S. cerevisiae*. Our results indicated that secretion improvements due to deletion of *cyt2* were applicable for this peroxidase as well, resulting in a 4.65-fold improvement in secretion with media supplementation (**Figure 2**). However, deletion of the O-glycosyltransferase *pmt2* had the strongest effect on secretion of PE-*vpl2*, improving secretion 9.45-fold with media supplementation and 2.68-fold without.

**Figure 1.**
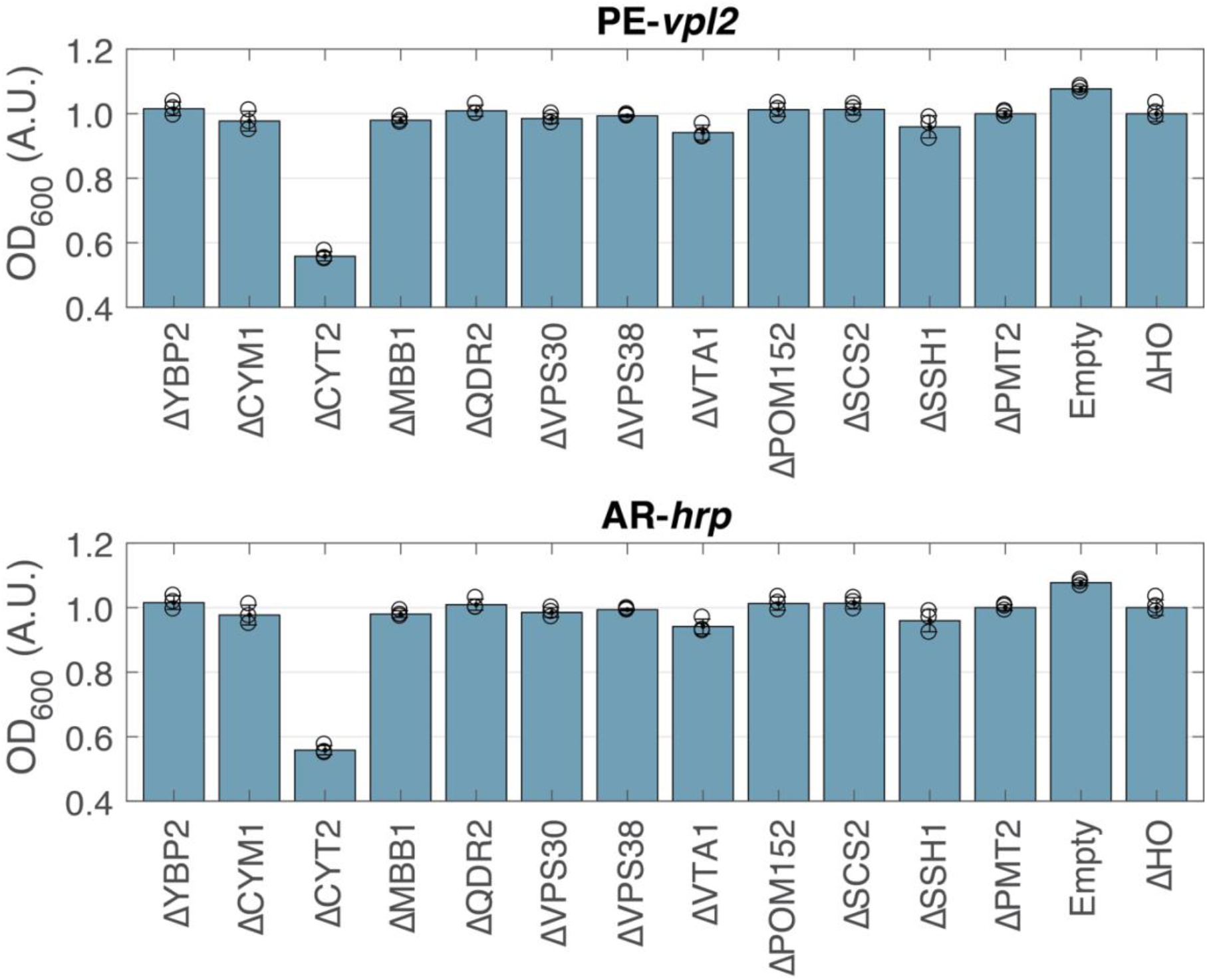
Culture growth measurements of selected mutants, normalized to ΔHO. The OD_600_ of culture supernatants of 12 BY4743 single knockout strains identified as having improved surface display of horseradish peroxidase (AR-*hrp*) were measured after 4 days cultivation at 20 C in YP media containing galactose and glycerol supplemented with calcium and hemin. Displayed values are normalized to OD_600_ of BY4743Δ*ho*. Bar graphs represent averages of three independent biological replicates and error bars represent one standard deviation.

**Figure 2.**
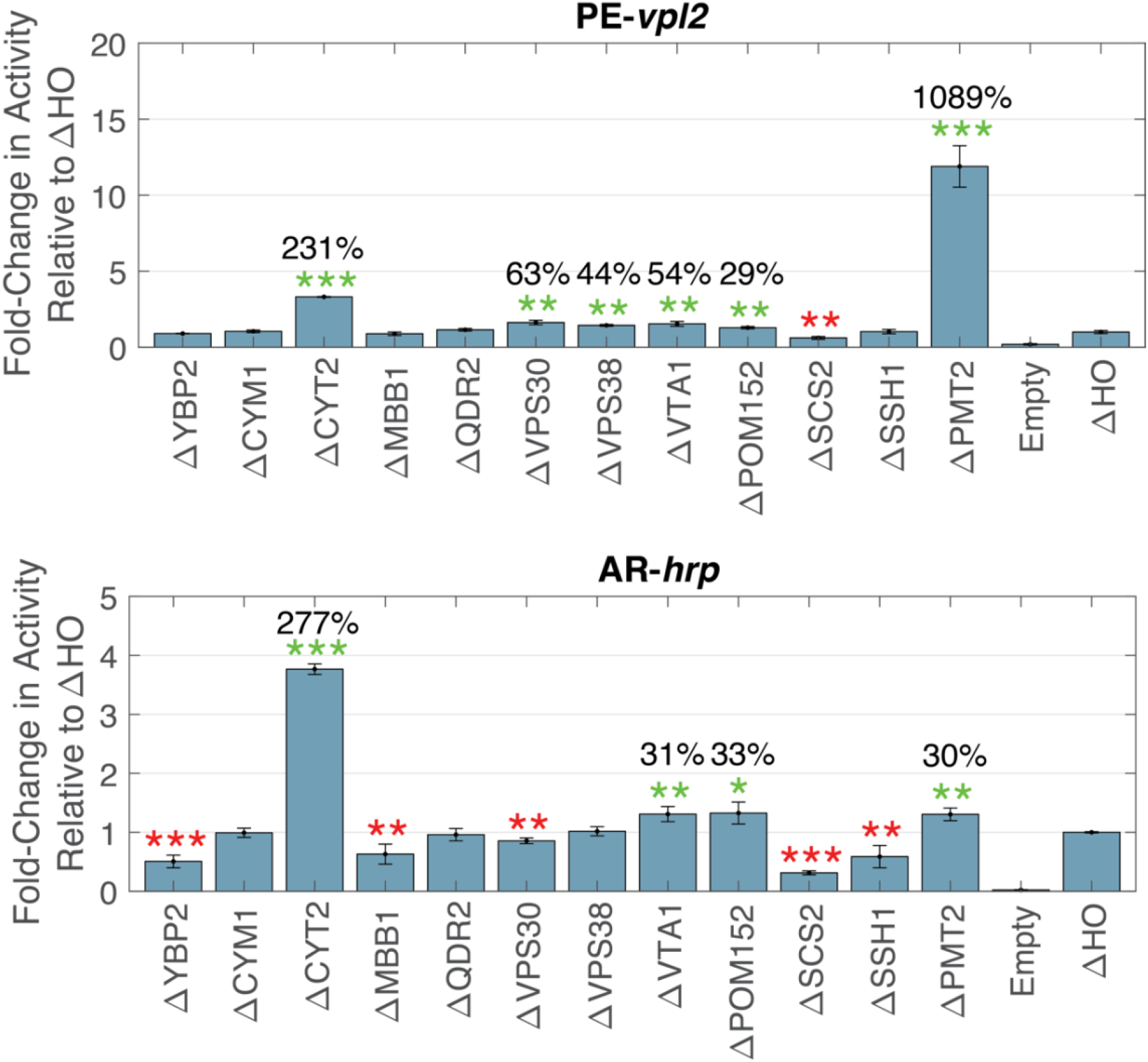
Extracellular activity levels in supplemented media of AR-*hrp* and PE-*vpl2* in selected mutants relative to ΔHO, corrected for OD_600_. Culture supernatants of 12 BY4743 single knockout strains identified as having improved surface display of horseradish peroxidase (AR-*hrp*) were assayed for ABTS activity as a proxy for secretion of PE-*vpl2* and AR-*hrp* after 4 days cultivation at 20 C in YP media containing galactose and glycerol supplemented with calcium and hemin. Activity levels are normalized to the OD600 of each culture and further normalized to activity levels of BY4743Δ*ho*. Bar graphs represent averages of three independent biological replicates and error bars represent one standard deviation.

Based on these results, heme cofactor incorporation and proper glycosylation appeared to be important yet sequential steps in the folding and processing of PE-*vpl2*. We hypothesized that the effects of deleting *cyt2* could be epistatic to those of deleting *pmt2* and constructed a double knockout BY4743Δ*cyt2*Δ*pmt2*. In attempt to help mitigate the growth defect caused by a homozygous *cyt2* knockout, we also constructed a double knockout heterozygous in the *cyt2* locus (BY4743Δ*cyt2*^-/+^Δ*pmt2*^-/-^). We observed normal growth in this heterozygous double knockout but surprisingly secretion of either peroxidase was reduced to that of wildtype both with and without media supplementation. The homozygous double knockout BY4743Δ*cyt2*Δ*pmt2* displayed growth defect but significantly outperformed either single knockout in secretion of either peroxidase, confirming our epistatic hypothesis (AR*-hrp*, 25.2-fold relative to wildtype; PE-*vpl2*, 57.0-fold; values after OD_600_ normalization) (**Figure 3**). Given the substantial improvements in secretion afforded by these two gene knockouts, we asked whether these effects were applicable to lignin-degrading peroxidases more broadly. We expanded secretion testing to include another versatile peroxidase from a different lignin-degrading fungus *Pleurotus ostreatus* (PO-*vp1*), a lignin peroxidase from *Gelatoporia subvermispora* (GS-*lip1*), and a manganese peroxidase from *Phanerochaete chrysosporium* (PC-*mnp1*). We observed almost undetectable levels of secretion in wildtype BY4743. Remarkably, deletion of *cyt2* and *pmt2* individually enabled detectable levels of secretion for all three lignin-degrading peroxidases (**Figure 3**). Moreover, the homozygous double mutant showed strong epistatic effects, particularly for PO-*vp1* (8.3-fold increase relative to Δ*cyt2* alone). These results indicate the BY4743Δ*cyt2*Δ*pmt2* significantly improves heterologous secretion of both class II fungal and class III plant heme peroxidases, and even enables secretion of lignin-degrading peroxidases having no detectable levels in wildtype BY4743.

**Figure 3.**
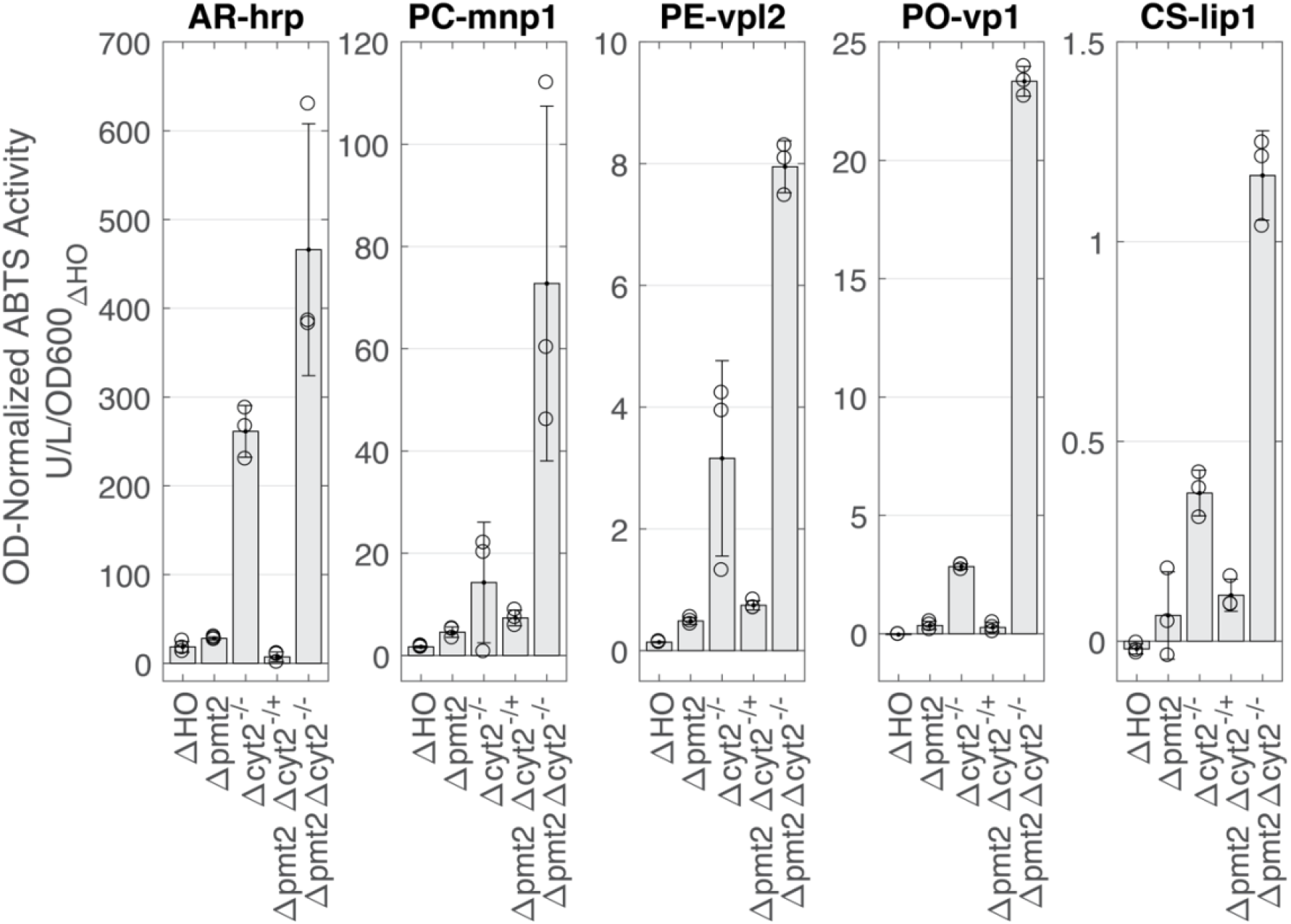
Epistatic effects of *pmt2* and *cyt2* mutants in the secretion of peroxidases. Culture supernatants of BY4743Δ*ho*, BY4743Δ*pmt2*, BY4743Δ*cyt2*, BY4743Δ*pmt2*Δ*cyt2*^*+/-*^, and BY4743Δ*pmt2*Δ*cyt2*^*-/-*^ strains expressing horseradish peroxidase (AR-*hrp*), *mnp1* from *P. chrysosporium* (PC-*mnp1*), *vpl2* from *P. eryngii* (PE-*vpl2*), *vp1* from *P. ostreatus* (PO-*vp1*), and *lip1* from *C. subvermispora* (CS-*lip1*) were assayed for ABTS activity after 4 days cultivation at 20 C in YP media containing galactose and glycerol supplemented with calcium and hemin. Activity levels are normalized to the OD_600_ of each culture and further normalized to activity levels of BY4743Δ*ho*. Bar graphs represent averages of three independent biological replicates and error bars represent one standard deviation.

We performed Western blotting of culture supernatants as well as whole cell extracts to better understand the effects of the gene knockouts on the molecular properties of PE-*vpl2* (**Figure 4**). Western blotting of whole cell extracts showed bands specific to PE-*vpl2* in all strains tested, indicating transcription and translation are not rate-limiting in the secretion of this peroxidase (**Figure 4, left**). BY4743Δ*ho* displayed bands at and above the expected unglycosylated molecular weight of PE-*vpl2* (36 kDa), with higher molecular weight bands presumably corresponding to increasingly glycosylated forms of the enzyme. Surprisingly, deletion of *pmt2* did not result in a noticeable effect on the molecular weight of the bands observed, although an increased amount of protein is evident. Deletion of *cyt2* resulted in a greater abundance of lower molecular weight bands, while the double deletion strain reflected a band pattern similar to the single *pmt2* deletion with greater protein levels as in the case of the single *cyt2* deletion. Bands at molecular weights lower than the expected size of the enzyme were also evident in strains deleted in *cyt2*, suggesting possible proteolysis. Blotting of extracellular protein revealed a single predominant band at molecular weight much higher than the dominant bands evident from intracellular blotting (**Figure 4, right**). This suggests that at steady state, the majority of the produced PE-*vpl2* is held up within the cell as evidenced by major bands at lower molecular weight, and only a heavily glycosylated form of the enzyme is secreted into the culture supernatant. Extracellular blotting also revealed that deletion of *pmt2* results in a decrease in molecular weight of the secreted enzyme glycoform, while deletion of *cyt2* enables increased secreted protein levels. Protein levels in extracellular blotting correlated well with observed activity towards ABTS in the context of single and double mutant testing focused on *pmt2* and *cyt2* (**Figure 3**), with the latter deletion displaying both highest activity levels as well as extracellular protein levels in the case of PE-*vpl2*. Differences in growth due to cultivation conditions for the slow-growing, respiratory-deficient *cyt2* strain may explain the difference in observed activity levels between screening of the 12 selected mutants (**Figure 2**) versus that of *pmt2* and *cyt2* mutants (**Figure 3**). Taken together, these data indicate that deletion of *pmt2* and *cyt2* result in increased secreted protein levels, resulting in greater observed extracellular ABTS activity levels, and that the deletion of both genes yields further improvements.

**Figure 4.**
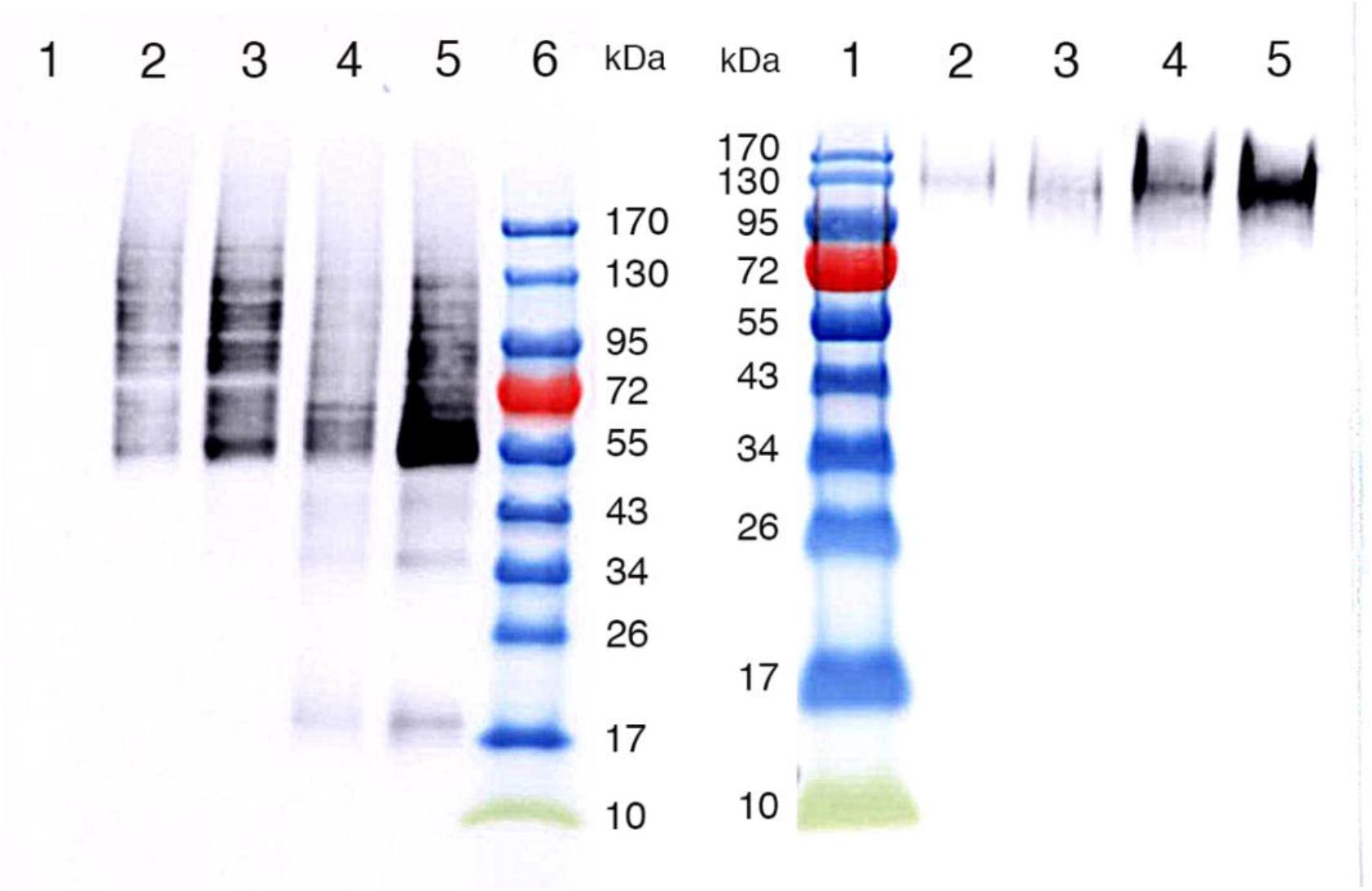
Western blotting of PE-*vpl2* produced by *pmt2* and *cyt2* mutants. **Left**: Whole cell protein extract blotting, with gel loading amounts normalized to cell density. Lanes: 1, empty vector control; 2, BY4743Δ*ho*; 3, BY4743Δ*pmt2*; 4, BY4743Δ*cyt2*; 5, BY4743Δ*pmt2*Δ*cyt2*; 6, ladder. **Right**: Extracellular protein extract blotting, with constant supernatant loading volume. Lanes: 1, ladder; 2, BY4743Δ*ho*; 3, BY4743Δ*pmt2*; 4, BY4743Δ*cyt2*; 5, BY4743Δ*pmt2*Δ*cyt2*. The expected molecular weight of PE-*vpl2* is about 36 kDa without glycosylation.

## Discussion

Secretion of heterologous proteins remains a frequently encountered and poorly understood challenge despite decades of research on secretory pathway components and the importance of heterologous protein production in multiple billion-dollar industries. We focused on a particularly challenging class of enzymes to produce heterologously, heme peroxidases from lignin-degrading fungi. Production of this enzyme class is the major bottleneck in the field of lignin degradation and the current inability to rapidly produce and test lignin-degrading heme peroxidases hampers the development of a biochemical route to woody biomass valorization as a renewable and abundant potential source of commodity chemicals. Previous studies have employed directed evolution to improve heterologous secretion of these enzymes in *S. cerevisiae*, but have relied on non-lignin-related substrates for screening^38,39^ and application of the evolved variants remains to be demonstrated on actual lignin substrates^40^. Given that the generally broad and unspecific substrate preferences of this class of enzymes remain poorly characterized for lignin-derived molecules, we opted for a strain engineering approach rather than protein engineering to improve heterologous secretion of these enzymes in *S. cerevisiae*. Since previous studies reveal that secretion improvements through random mutagenesis of yeast genes arise mainly from loss-of-function mutations^19^, we screened a barcoded deletion collection^34^ of single null mutants of non-essential genes in *S. cerevisiae*, which revealed that deletion of *pmt2* and *cyt2* resulted in substantial increases in heterologous heme peroxidase secretion for both a model plant peroxidase as well as a diverse set of fungal lignin-degrading peroxidases.

The improvement in secretion due to deletion of *cyt2*, cytochrome c heme lyase, reveals heme biosynthesis and/or trafficking as a limiting factor. *S. cerevisiae* does not have any known transporters specialized for exogeneous heme, yet media supplementation with free hemin was shown to be beneficial in this study. In the production of cytochromes P450, heme biosynthesis has been shown to be an engineering target for protein yield improvements^41^ and previous studies have achieved improved peroxidase production through supplementation with heme precursors and upregulation of the heme biosynthetic pathway^42^. *Cyt2* however is not directly involved in heme biosynthesis but utilizes heme for maturation of cytochrome c1^43^. This suggests that reducing heme demand through deletion of *cyt2* yields more available heme for the production of heterologous heme peroxidases. Greater heme availability may also allow greater production of native peroxide-detoxifying enzymes such as cytochrome c peroxidase and catalase^44^, mitigating possible oxidative effects of heterologous peroxidase expression in the secretory pathway. This deletion however results in an inability for the strain to grow on non-fermentable carbon sources, meaning that while cellular productivity is increased, cell density under inducing, protein-producing conditions is limited. Nevertheless, this result indicates that *cyt2* presents an important candidate for further strain engineering efforts. For example, complementation by *cyt2* homologs or evolution of *cyt2* variants having reduced function would mitigate growth defects while increasing the supply of heme available for heterologous proteins. Moreover, factors involving the transport of heme into the secretory pathway remain unknown and could further enhance peroxidase secretion in conjunction with *cyt2* expression modulation.

Our screen revealed that deletion of *pmt2* had the greatest positive impact on secretion of fungal lignin-degrading peroxidases of the mutants tested. This effect was notably observed for multiple members from different lignin-degrading peroxidase classes from a variety of fungal species, suggesting that a general feature of this enzyme class results in misprocessing in yeast. *Pmt2* is a member of the family of O-mannosyltransferases involved in ER protein quality control. Fungal lignin-degrading peroxidases are known to have both N- and O-linked forms of glycosylation of the simple mannose type. Western blotting of intercellular as well as extracellular protein fractions revealed bands presumably corresponding to glycoforms of PE-*vpl2* at very high molecular weight (**Figure 5**). Hyperglycosylation is a previously characterized challenge in secretion of heterologous proteins in *S. cerevisiae*^12,29^, including that of horseradish peroxidase^29^. Repeated mannosyl transfer would yield a sterically hindered protein that may pose greater difficulty in export through the yeast peptidoglycan cell wall matrix. Why *pmt2* appears to target fungal lignin-degrading peroxidases in particular for excessive glycosylation is unclear. Despite their structural and glycosylation similarities, the effect of *pmt2* deletion was significantly greater for this class enzymes compared to horseradish peroxidase. This suggests protein sequence determinants for misprocessing by *pmt2*. As an ER protein quality control component, targeting by *pmt2* also seems to indicate protein misfolding issues. Incubation of PE-*vpl2*-expressing cells in the presence of colorimetric peroxidase substrates (*e*.*g*. ABTS) gave no color change. As these substrates could be expected to diffuse into the yeast cell wall, this suggests that the enzyme is either trapped but inactive within the cell wall matrix, or otherwise does not reach the cell periphery.

The combination of *cyt2* and *pmt2* deletions revealed synergistic effects on secretion of all peroxidases tested. This study demonstrates that single null mutant screening affords significant improvements in heterologous protein production and more importantly serves as a proof-of-concept to enable subsequent screens focusing more on epistatic effects of gene expression modulation. Deletion and/or overexpression of other native yeast genes in the context of the *cyt2/pmt2* deletion strain background would almost certainly yield further secretory improvements, which may have had only modest effects in the context of the wildtype strain background. Variable gene expression modulation using, for example, plasmid-based CRISPRi/a libraries would greatly expand the scope by enabling screening of essential yeast genes, facilitating rapid testing of library member combinations, and identify optimal levels of gene expression. While this study searched for components native to yeast that prevented successful production of lignin-degrading peroxidases, it may be that yeast lack components present in lignin-degrading fungi required for proper folding and processing of these enzymes. These may be identified for example by screening strains co-expressing genomic cDNA libraries from lignin-degrading fungi using the same strategy described in this study. Finally, we anticipate this study to serve as the first step towards directed evolution of cellular components rather than of proteins of interest to improve heterologous production. The genomic targets identified here can be mutagenized and screened for variants with greater secretion enhancement than by deletion of those targets alone while minimizing impacts on cell growth. Methods involving *in vivo* mutagenesis and continuous selection currently being developed would further accelerate this approach. The results of these efforts would help augment our understanding of factors implicated in heterologous protein production, expanding our ability to reliably and efficiently produce valuable protein-based therapeutics and catalysts at scale.

## Methods

The homozygous diploid yeast deletion collection was kindly provided in pooled format by the Stanford Genome Technology Center and propagated as previously described in YPD. Transformation of the peroxidase-surface display fusion vector pL158A-HRP was carried out by the lithium acetate method and plated on synthetic defined media plates lacking leucine to select for transformants. Transformants were pooled from 8 plates into YPD + 15%*w/v* glycerol giving at least 36000 transformants for pL158A-HRP and were frozen at -80 C.

27 μl of library stock was used to start a 2 ml non-inducing, non-repressing synthetic defined medium containing 2% raffinose and 1% sucrose lacking leucine for auxotrophic selection. After 20 hours growth (30 C, 300 rpm), 100 ul culture was diluted into 2.9 ml synthetic defined medium containing 4% galactose and lacking leucine. After 16 hours induction (20 C, 300 rpm), 10 million cells were harvested in quadruplicate and labeled as previously described^31^. Briefly, cells were washed twice with 1 ml buffer (PBS + 0.1% BSA) before being resuspended in biotinyl tyramide reaction buffer (100 μM biotinyl tyramide, 100 μM hydrogen peroxide, PBS + 0.1% BSA, pH 7.4) at cell density of 10^6^ per ml in 15-ml conical centrifuge tube. Cells were labeled statically on ice for 5 minutes before quenching with 0.5 ml buffer containing Trolox (10 mM Trolox in PBS + 0.1% BSA). Cells were pelleted into a 1.5-ml microcentrifuge tube and washed twice with buffer. Samples were labelled using a 1/50 dilution of anti-Myc-AlexaFluor647 conjugate (Cell Signaling Technologies) and a 1/10 dilution of streptavidin-phycoerythin conjugate (Jackson Laboratories) in 200 μl buffer for 1.75 hours at 4 C in the dark with rotation. Samples were washed twice with 200 μl buffer and immediately use for cell sorting.

Fluorescence-assisted cell sorting was performed on a Sony SH800 cell sorter with compensation for PE and AlexaFluor647 channels. 1.2 to 1.4 million events were sorted in total for the four replicates, collecting cells in 3 ml synthetic defined medium containing 2% glucose and lacking leucine. “Unsorted” control samples were prepared by collecting all events into a single culture. Genomic DNA was extracted using a YeaStar Genomic DNA kit (Zymo Research) after 21 hours outgrowth (30 C, 300 rpm). “Non-displaying”, “displaying”, and “unsorted” samples showed comparable final cell densities. Cultures were backdiluted into inducing growth media as above after 26 hours outgrowth to validate sorting. Biotinyl tyramide labeling was performed as above, and fluorescent labeling was performed at 1/100 and 1/50 dilutions, respectively.

2 μl of genomic DNA was used to amplify strain barcodes by PCR (Q5 NEB master mix, 22 cycles) using primers containing sequence-optimized spacers to maximize nucleotide diversity in Illumina sequencing. DNA amplification was performed with two different spacer primers as technical duplicates to minimize PCR amplification bias due to primer sequence. PCR products were gel-purified and used for a second round of PCR amplification (Q5 NEB master mix, 7 cycles) using custom primers to attach Illumina read sequences. PCR products were gel purified and their concentration quantified using a Qubit. Products were pooled and sequenced using an Illumina HiSeq 4000 2×75bp. Strain-barcode matching and counting were performed using a Levenshtein distance of 2 and neglecting barcodes appearing only once. Statistical analysis was performed using the *edgeR* software package in R, taking raw count data from replicate sort samples as data input. An adjusted counts per million of more than 10 in at least 6 samples was used to filter out strains with low abundance. Log fold change of strain counts between “displaying” and “non-displaying” samples were used to assess gene deletion effect on secretion. Individual single null mutant strains *pom152, ybp2, cym1, vps30, cyt2, pmt2, qdr2, vps38, ssh1, scs2, ssb1 and ho* were kindly provided by Angela M. Chu of the Stanford Genome Technology Center and propagated on YPD plates. Double knockout strains were constructed by homologous recombination in the *pmt2* mutant background using his3 and ura3 as selectable markers to replace *cyt2*. Transformation of pL158A vectors was performed using the lithium acetate method with selection on synthetic defined plates lacking leucine. Transformants were picked into 96-well plates with synthetic defined growth medium containing 2% glucose and lacking leucine and grown overnight (24 hours, 30 C, 450 rpm). Cultures were diluted 1/10 into YP medium containing 2% galactose and 3% glycerol and induced for 4 days (20 C, 450 rpm). Peroxidase activity was assayed using a Synergy HTX plate reader at 25 C using 10 mM ABTS, 100 μM hydrogen peroxide, and one of the following: 50 mM potassium phosphate, pH 6.0, for horseradish peroxidase; 50 mM sodium tartrate, pH 3.5, for versatile and lignin peroxidases; and 50 mM sodium malonate, pH 4.5, and 1 mM MnSO_4_ for manganese peroxidases.

